# Validation of *de novo* designed water-soluble and transmembrane proteins by *in silico* folding and melting

**DOI:** 10.1101/2023.06.06.543955

**Authors:** Alvaro Martin Hermosilla, Carolin Berner, Sergey Ovchinnikov, Anastassia A. Vorobieva

## Abstract

*In silico* validation of *de novo* designed proteins with deep learning (DL)-based structure prediction algorithms has become mainstream. However, formal evidence of the relationship between a high-quality predicted model and the chance of experimental success is lacking. We used experimentally characterized *de novo* designs to show that AlphaFold2 and ESMFold excel at different tasks. ESMFold can efficiently identify designs generated based on high-quality (designable) backbones. However, only AlphaFold2 can predict which sequences have the best chance of experimentally folding among similar designs. We show that ESMFold can generate high-quality structures from just a few predicted contacts and introduce a new approach based on incremental perturbation of the prediction (“*in silico* melting”), which can reveal differences in the presence of favorable contacts between designs. This study provides a new insight on DL-based structure prediction models explainability and on how they could be leveraged for the design of increasingly complex proteins; in particular membrane proteins which have historically lacked basic *in silico* validation tools.

## Introduction

*De novo* protein design^1,2^ is the generation of protein sequences and structure from scratch with no detectable homology to natural proteins of similar structure^3–5^ and functions^6,7^. Most *de novo* protein design pipelines build on a common paradigm: (i) sequence-free protein backbones are assembled with selected structure properties and (ii) used to guide combinatorial design of sequences compatible with the desired folded state^1,8^. Sequences are then experimentally evaluated for expression and for folding into the designed structure. The percentage of designs that meet the experimental criteria for success is largely variable. Entire batches of designs can fail to express or to fold because they were generated based on non-designable protein backbones. *In silico* generated protein backbones are considered “designable” if they incorporate key structural features of natural proteins and can encode amino acid sequences with real global minima^7,9^. In practice, there are few explicit guidelines to help generate designable backbones, and the task requires a significant level of expertise. Even when designable backbones are available, the *de novo* design of certain classes of proteins, such as complex β-sheet folds^7,10,11^ or membrane proteins^12,13^, can be challenging because the design algorithms or energy functions can fail to capture structural or biophysical properties specific to these folds. As a result, the success rate of *de novo* design ranges from 50% (mostly α-helical folds^14^) to less than 5% for some of these challenging cases.

To improve the success rate, the designs are often filtered and ranked for experimental validation by predicting *in silico* the most likely 3D structure adopted by the linear chain of amino acids (*ab initio*) to probe the sequence-structure compatibility^15,16^. However, classic *ab initio* predictions (e.g. Rosetta^17^) often lead to inconclusive results as the complexity of the protein fold increases with size, contact order and β-sheet content^18^. For membrane proteins, the implementation of an implicit membrane environment^19,20^ makes it possible to screen only relatively simple designs such as self-assembling transmembrane α-helical peptides^21–23^. As a result, membrane protein (MP) design still lacks many *in silico* validation tools available for water-soluble proteins^13^.

Recently, deep learning (DL) models such as AlphaFold2^24^, RoseTTaFold^25^, and language models (LM) (e.g ESMFold^26^, OmegaFold^27^ or RGN2^28^) have out-performed classic *ab initio* folding simulations. AlphaFold2, RoseTTAFold and ESMFold networks have also been inverted to design new protein sequences^29–31^. We sought to provide a detailed and quantitative overview of the opportunities and limitations of using AlphaFold2 and protein language models – exemplified here by ESMFold - for *in silico* validation and ranking of *de novo* designed proteins. We assembled three datasets of experimentally characterized synthetic proteins considered as “challenging targets” for classic *ab initio* structure prediction: (i) sequences generated based on un-designable backbones, (ii) β-sheet proteins designed to fold in water and (iii) proteins folding in lipid membranes. We predicted the structures of the proteins in the datasets using standard and modified AlphaFold2 and ESMFold pipelines to identify conditions and metrics that could reveal differences between the sequences that successfully passed or failed experimental validation.

### Designable and un-designable backbones

We first sought to assess AlphaFold2 and ESMFold capacity to distinguish designs generated based on designable and un-designable backbones. We assembled two datasets of eighty-three sequences designed to fold into β-barrel structures^7^. β-barrels are formed from one unique β-sheet that curves on itself to close into a tube-shaped structure. A set of forty-two designs incorporated specific structure features (kinks in the protein backbones) that helped release strain in the curved β-sheet, thereby stabilizing the desired folded state and achieving a success rate of about 35% (“designable” dataset). The second dataset contained forty-one designs generated with perfectly regular β-strands lacking structural kinks (“un-designable” sub-group)^32,33^. None of these sequences folded into the expected monomeric structure. Instead, they failed to express or were insoluble. All eighty-three designs (designable and un-designable) passed the classic *ab initio* folding test and showed a funnel-shaped *in silico* folding landscape converging to the design model (Figure S1). Hence, classic *ab initio* structure prediction was unable to distinguish designs from the two datasets. We modeled the 3D structures with standard AlphaFold2 and ESMFold pipelines and used two metrics to assess model quality: (1) the confidence assigned to the prediction (predicted local distance difference test, pLDDT) and (2) the prediction accuracy, calculated as the root mean square deviation (RMSD) of the predicted structure to the design model.

Both ESMFold and AlphaFold2 predicted the structures of *de novo* designed proteins from a single sequence (without a multiple sequence alignment input), which clearly clustered into distinct “designable” and “un-designable” populations (Figure 1a,b). “Designable” sequences yielded, in average, similar structures to the expected design models and which were attributed higher confidence (Figure 1c). EMSFold performed significantly better at distinguishing the design populations in the tested dataset; when the pLDDT score was higher than 0.75 and the RMSD between the predicted structure and the design model lower than 2.0 A, the chance to get a “designable” β-barrel backbone was 100%.

**Figure 1:**
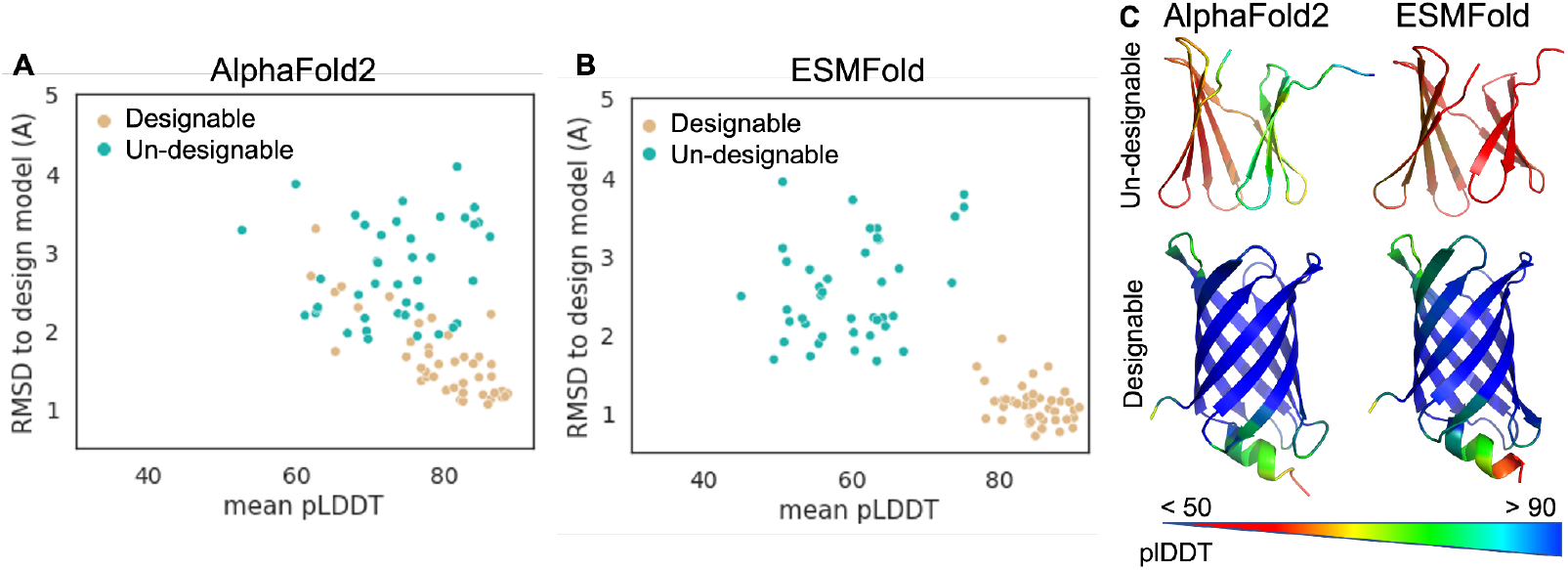
AlphaFold2 and ESMFold predict backbone designability of de novo designed proteins. AlphaFold2 (a) and ESMFold (b) predictions on sequences from the designable and un-designable sets sample different pLDDT and RMSD values (c) Examples of AlphaFold2 and ESMFold predictions colored based on local pLDDT.

### Folded and misfolded sequences from similar design sets

Even with designable backbones as starting points, the probability of success of *de novo* design ranges from 5 to 50 % within sets of designs typically featuring relatively similar computed energies and sequences (40-75 % sequence identity). The determinants of success/failure remain elusive and likely involve more subtle differences than those underlying designable backbone properties. We sought to determine whether these differences were captured by AlphaFold2 and ESMFold.

#### Overview of the datasets

We assembled datasets of *de novo* designed β-sheet proteins folding in water (water-soluble) and in lipid membranes (transmembrane) and labeled according to their experimental behavior: expressed, soluble (in water or detergent micelles) and successfully folded. All-β-sheet folds were selected here because they represent particularly challenging targets for classic *ab inito* folding simulations due to the presence of long-range (and potentially ambiguous) interactions between β-strands. Despite sharing similar structure properties, water-soluble and transmembrane β-sheet folds have very different sequence properties^34^. The water-soluble dataset comprised forty-two β-barrels generated based on designable backbones. Fifteen of these sequences were correctly folded (∼35 % success rate) as attested by the presence of a detectable monomeric species with the expected β-sheet secondary structure. No difference between these fifteen folded designs and the twenty-seven “failed” (non-expressing, aggregated or misfolded) designs was detected using Rosetta *ab initio* structure prediction. The transmembrane dataset is, to our knowledge, the largest published dataset of experimentally characterized synthetic membrane proteins, composed of 115 sequences designed to fold into small 8-strands transmembrane β-barrels (TMB)^35^. A major challenge for TMB design is the necessity to incorporate local sequence/structure frustration to modulate the folding kinetics of the polypeptide chain. All twenty-five designs in our dataset that did not incorporate frustrated residues (strong β-sheet designs) failed to express in *E. coli*^35^. Among the ninety designs generated with local secondary structure frustration (weak β-sheet designs), seventeen folded as expected into monomeric β-sheet proteins in detergent micelles (∼19 % success rate)^35^. TMB designs were not evaluated *in silico* due to technical limitations of *ab initio* folding simulations^20,36^.

#### Sequence-structure compatibility assessment with AlphaFold2 folding

The capacity of AlphaFold2 to predict the structure of *de novo* designed water-soluble proteins from a single sequence has been demonstrated on several occasions^26,37–39^. Here, we looked for differences between models generated for experimentally folded and misfolded sequences. We found that the two groups of designs sampled significantly different distributions of pLDDT and RMSD (p-values 10^−11^ and 10^−5^, respectively) (Figure 2a,b). Predicted structures of folded designs were, on average, attributed higher pLDDT confidence and were more similar to the respective design models than the designs that misfolded (Figure 2b,c). Out of five generated models, successfully folded designs yielded significantly more high-quality models (pLDDT >= 0.85 and RMSD <= 1.75) than misfolded designs (average of 3.0 and < 1.0, respectively) (Figure S2a). Considering the lowest-RMSD model per input (Figure S2b), the chance of picking a “folding” sequence was 90 % when pLDDT was higher than 0.85 and RMSD lower than 1.75 (with a false negative rate of 21 %). The Receiver Operator Characteristic (ROC) curves are shown in Figure S2c.

**Figure 2:**
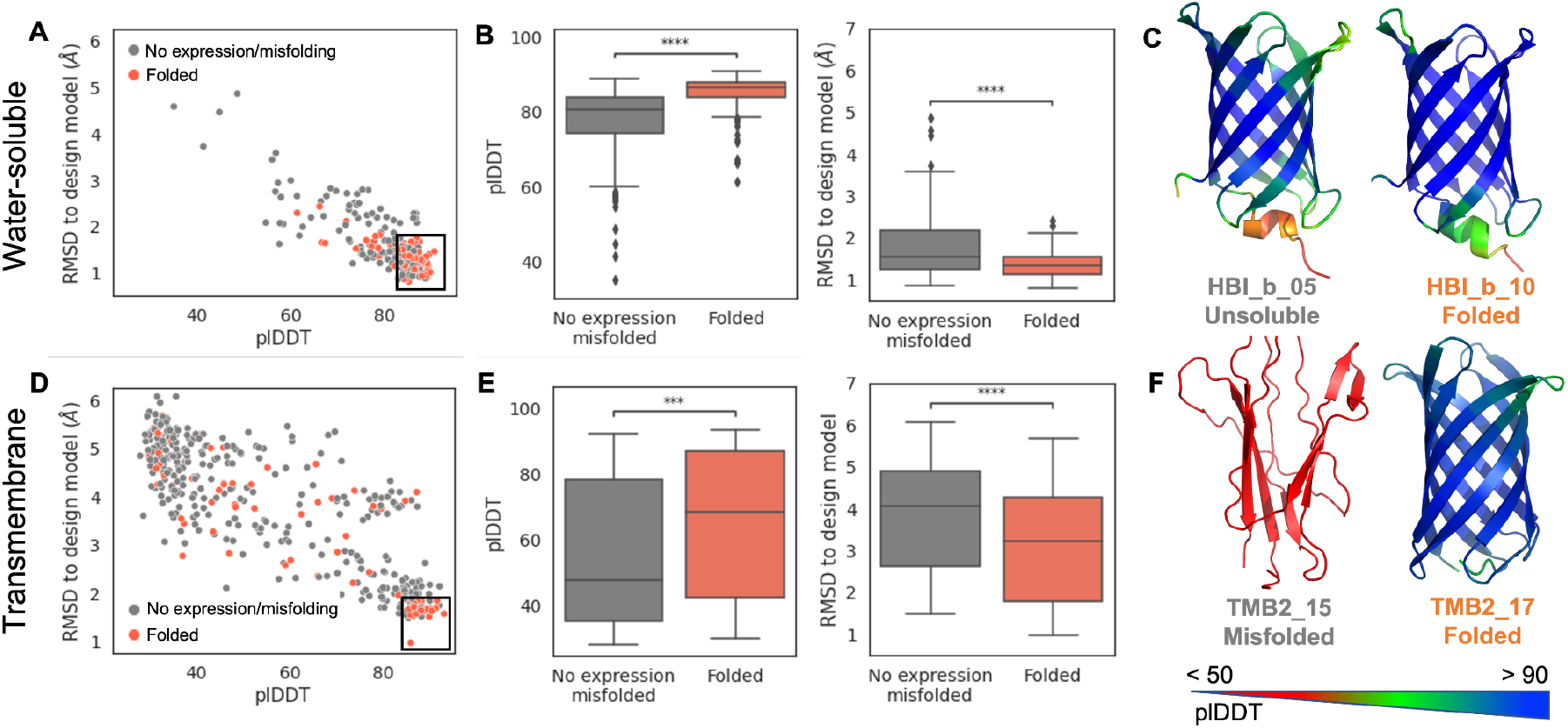
AlphaFold2 generates significantly different predictions for successfully folding and misfolding water-soluble (a-c) and transmembrane (d-f) β-barrel designs.

In contrast to water-soluble proteins, the capacity of AlphaFold2 to predict the structure of designed membrane proteins from a single sequence was unknown (MPs represent less than 2 % of structures deposited in the PDB and hence of AlphaFold2’s training dataset^40^). We evaluated “default” settings AlphaFold2 on our dataset of 115 *de novo* designed TMBs. Surprisingly, the designs that failed to express experimentally yielded predictions with the best pLDDT and RMSD scores (p-value(pLDDT) = 10^−4^; p-value(RMSD) = 10^−4^) (Figure S3a,b). We reasoned that the “strong β-sheet” designs in our dataset (which largely failed to express) might also feature more distinct 3D structure signatures than the frustrated “weak β-sheet” designs and therefore yield higher quality predictions uncorrelated to experimental success (Figure S3c). We decided to focus our analysis on the sequence space that can be expressed and experimentally evaluated and filtered-out “strong β-sheet” sequences based on empirical β-sheet (50%, RaptorX^41^) and aggregation propensity cutoffs (1750, Tango^42^) (Figure S3d). However, we found no difference between the folded and misfolded designs (pvalue(pLDDT) = 0.99; pvalue(RMSD) = 0.33) among the retained eighty-one sequences. We hypothesized that “weak β-sheet” TMBs sequences were challenging targets for single sequence AlphaFold2 because of the designed sequence-structure frustration, and that increasing the number of recycles through the Evoformer^24^ could help resolving weak 3D contacts. The predictions improved with forty-eight recycles (instead of three): successfully folded designs sampled significantly better distributions of pLDDT (pvalue = 10^−4^) and RMSD (pvalue = 2*10^−4^) than the designs that misfolded (Figure 2d,f). AlphaFold2 generated at least one full β-barrel structure for thirteen out of sixteen folding designs, while misfolding sequences yielded mostly low confidence, unstructured, models (Figure S2d, Figure 2f). Considering the lowest-RMSD model per input (Figure S2e), the chance of getting a folding sequence was 48 % when pLDDT was higher than 0.85 and RMSD lower than 1.85 (see the Receiver Operator Characteristic (ROC) curves in Figure S2f). This represents a 2.5-times increase of the success rate by comparison to the “naive” design selection. Hence, a carefully designed AlphaFold2 screen taking into consideration the biophysical specificities of the (water-soluble or membrane) fold could improve the success of *de novo* protein design by 2-3 times.

#### Sequence-structure compatibility determination with ESMFold

ESMFold is a large protein language model trained to decode a protein structure from a single sequence of amino acids. The software comprises a language module (LM) predicting residue contacts from the sequence input, and a structure module (SM) generating 3D coordinates based on the sequence and the predicted contact^43^. Protein language models such as ESM have been developed to capture protein properties from a single sequence. Therefore, ESMFold is often considered as a suitable tool for the analysis of synthetic (*de novo* designed) and orphan protein sequences, for which few homologs can be found. Strikingly, it generated high pLDDT, low RMSD models for nearly all sequences tested here (Figure 3). It could predict the structures of *de novo* designed TMBs with comparable performance to water-soluble β-barrels and made almost no distinction between experimentally folded and misfolded designs (Figure 3b-d and ROC curves in Figure S4a-b). The predicted TMscore (pTM) score had only marginally higher predictive power in case of TMB designs over the pLDDT metric (Figure S4).

**Figure 3:**
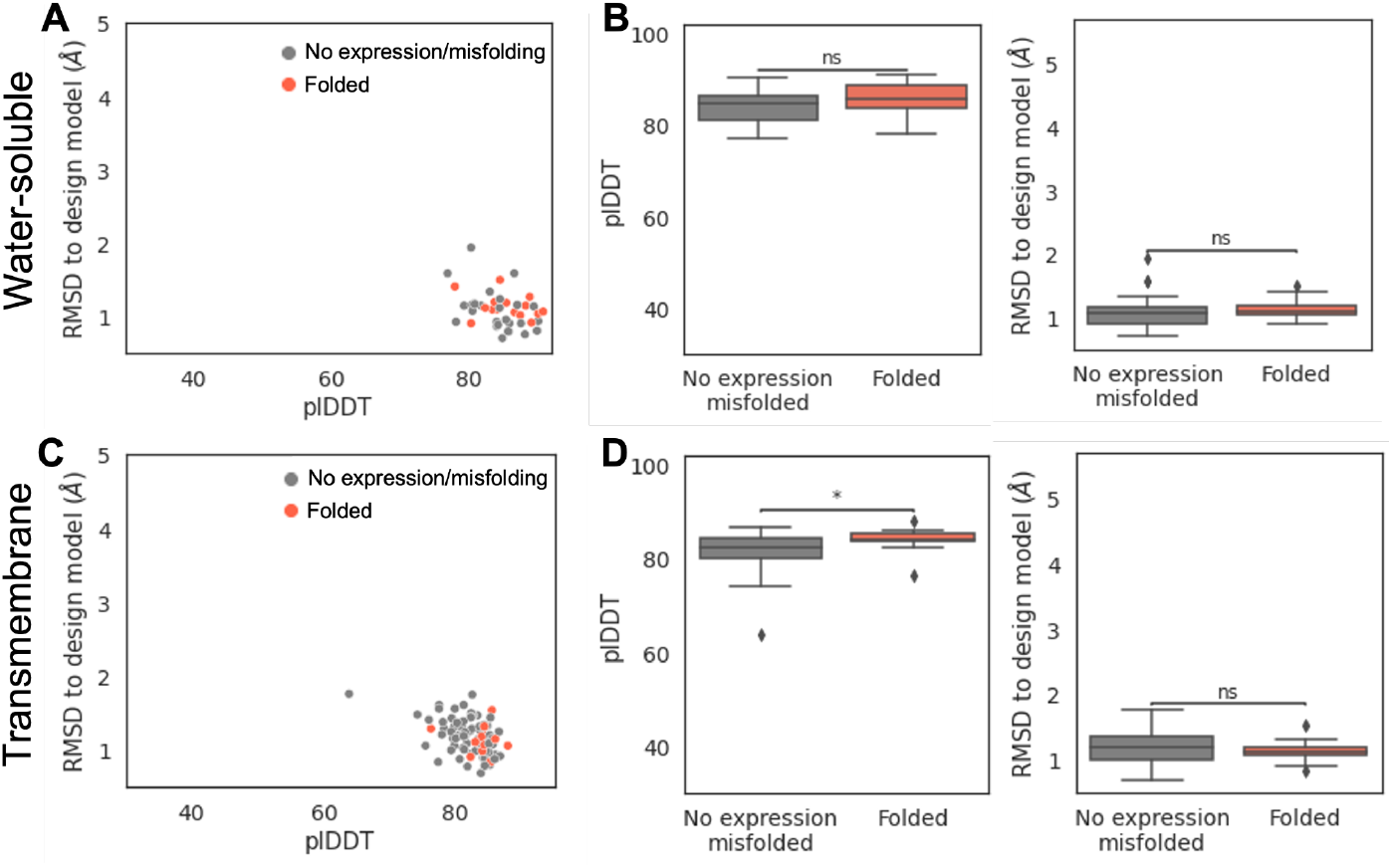
ESMFold generates high-confidence predictions closely resembling the design models for water-soluble (A-B) and transmembrane β-barrels (C-D).

#### Model extrapolation to novel membrane protein designs

To assess the capacity of AlphaFold2 and ESMFold to extrapolate beyond previously published designs and natural β-barrels structures, we tested a set of forty (including seven successfully folded) unpublished sequences designed to fold into 12-stranded TMBs without homology to natural proteins and with β-barrel architectures not yet represented in the Protein DataBank (outside diameters 21.5 - 25.8 Å). Overall, the performance of the two models was similar on the new and the previously tested datasets. While we observed significantly better performance of AlphaFold2 on experimentally folding over misfolding sequences (Figure 4a,b), ESMFold generated high-confidence structures closely resembling the design models for almost all tested designs, irrespective of experimental outcome (Figure 4c). Hence, AlphaFold2 and ESMFold were able to generate accurate single-sequence predictions of complex membrane proteins without sequence nor structure similarity to natural proteins. However, only AlphaFold2 was able to distinguish properly folded from misfolded design. Although ESMFold generated accurate structures for most designs in the new dataset, two predictions differed from the design model at the level of the connectivity between the first and last β-strands. To investigate the origin of the discrepancy, we developed an ESMFold Colab notebook that reports the residue contacts predicted by the LM and the SM. We used the notebook to predict the structures of one 8-stranded and three 12-stranded TMBs, with outside diameters ranging from 16.0 to 25.8 Å. We found that the contribution of the LM to long-range contacts detected by the SM (*TopL/5 LM/SM contribution*) decreased when the diameter of the β-barrel increased (Figure 4d). No contacts were predicted between the first and the last β-strands of the largest TMB. Yet, a closed β-barrel with the expected β-strands connectivity was generated by the SM. The observed cooperation between the LM and SM might explain the consistent results achieved by ESMFold on *de novo* designed sequences.

**Figure 4:**
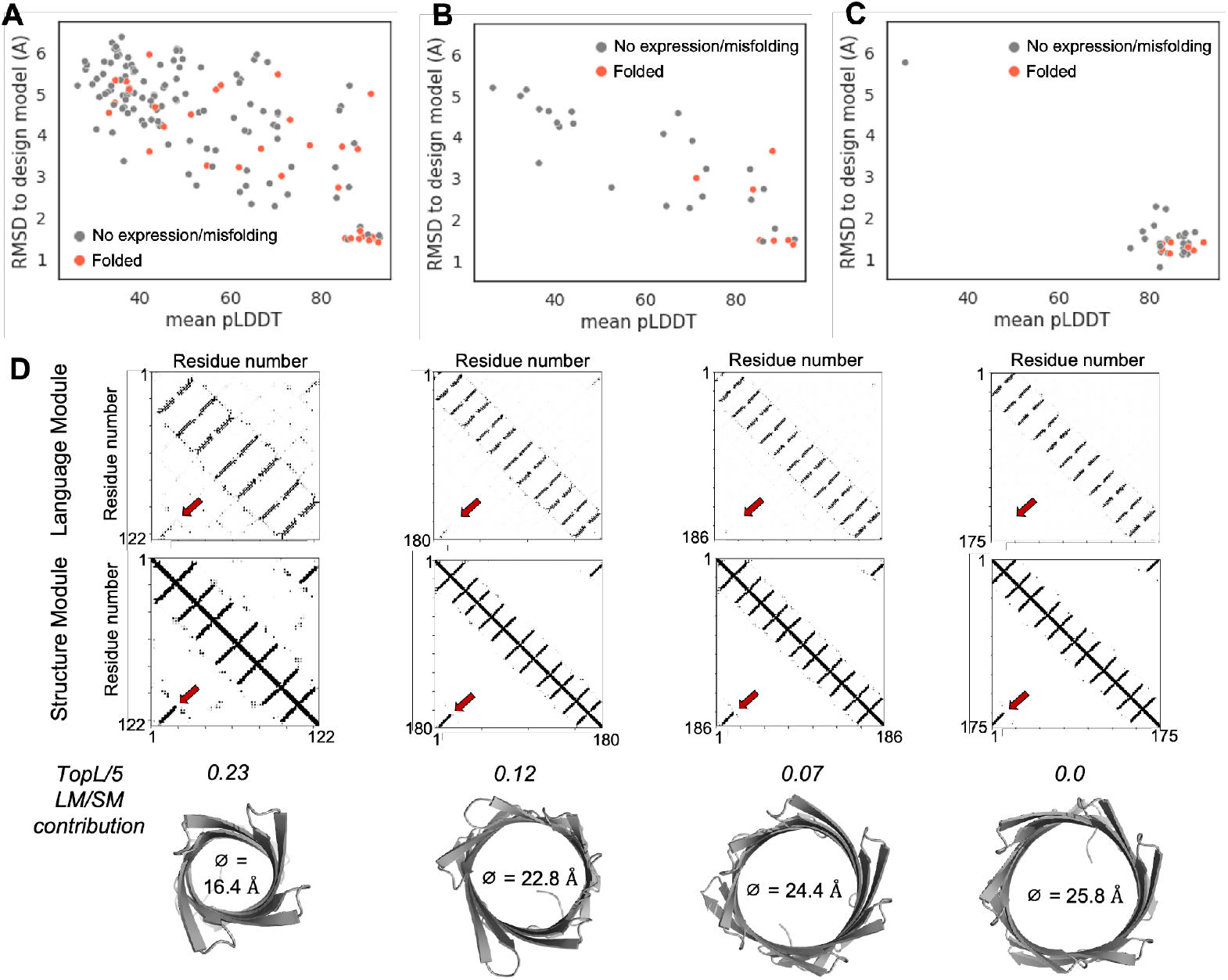
AlphaFold2 can distinguish successfully folding from failed (misfolding or non-expressing) TMB designs with new-to-nature structures and sequences; using all five models (a) or the top-ranked RMSD model (b). (c) ESMFold generates high-confidence, low RMSD, predictions for nearly all novel TMB sequences. (d) Fewer long-range contacts (TopL/5) are predicted by the LM as the diameter of the β-barrel increases. Yet, the TopL/5 contacts are correctly predicted by the SM.

#### In silico melting reveals difference between folding and misfolding designs

We investigated the robustness of the ESMFold prediction and the rescue capacity of the SM as a function of the input sequence by randomly perturbing the LM. We implemented a new approach in a Colab notebook, dubbed *in silico* melting, to systematically sample predictions with incremental levels of noise. To introduce noise into the prediction, we took advantage of the masked token built-in to train the ESM language model^26^. The prediction was noised by masking a subset of randomly selected residues in the input sequence. For each level of noise, a success rate is calculated based on the number of sampled predictions with a pTM score of 0.75 or higher. Four *de novo* designed proteins with a high-resolution structure in the PDB were initially analyzed. The predictions obtained for the α-helical bundle^14^ and the α/β TOP7^3^ designs were surprisingly robust; accurate structures were generated despite masking 90-95 % of the sequence. By contrast, the success rate dropped at around 55 % noise for the water-soluble^7^ and the transmembrane β-barrels^35^ (Figure 5a). Hence, the SM can predict α-helices rich structures using just a few LM-predicted contacts but requires more contacts information for β-sheet proteins.

**Figure 5:**
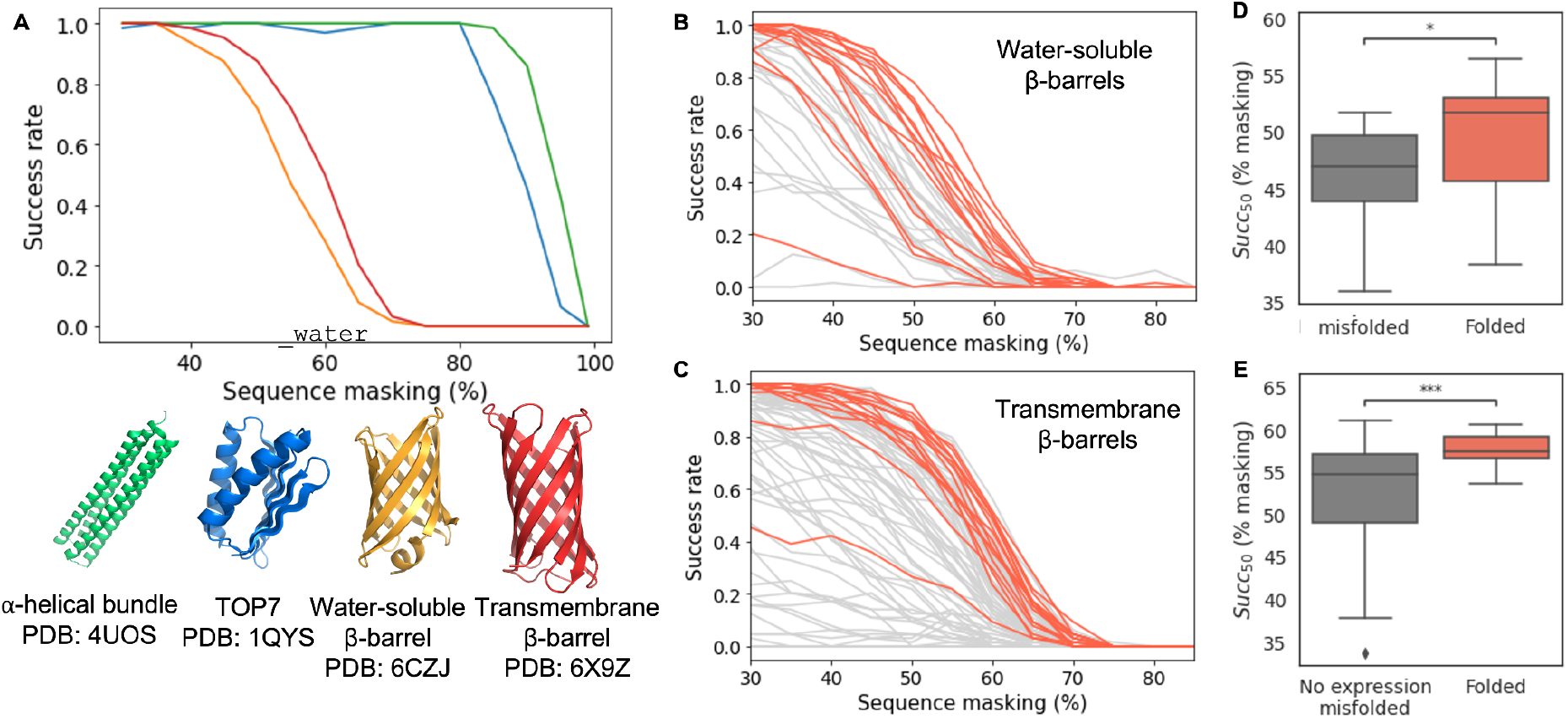
(a) ESMFold in silico ‘melting’ curves for four de novo designed proteins. (B-C) Predicted robustness for designs from the water-soluble (b) and transmembrane (c) β-barrels datasets. Folded designs are shown as red curves. (d-e) Distributions of sigmoidal inflection point values for water-soluble (d) and transmembrane (e) β-barrels. P-values of 2*10^−2^ and 4*10^−4^, respectively

We next used the *in silico* melting to compare designs with similar structures and sequences in the water-soluble and transmembrane β-barrel datasets. In both datasets, ESMFold predictions for successfully folded designs were more robust to sequence masking than the designs that failed to fold (Figure 5b-c). We fitted a sigmoidal model to the simulated melting curves to extract the inflection points of predictions (*Succ*_*50*_) and found a significant difference between the folded and misfolded designs (Figure 5d-e). While all sequences yielded similar final EMSFold predictions, sequence pattern masking to probe the robustness of structure contacts might provide a new insight into the properties of generated designs.

## Discussion

AlphaFold2 and ESMFold are routinely used for the *a*ssessment of *de novo* designed protein sequences and to compare the performance of design methods. However, the relationship between a good prediction and the expected experimental outcome has not been formally demonstrated. We use experimentally characterized sets of *de novo* proteins - designed to fold in water or lipid membranes - to test the models. Our results support previous observations that both AlphaFold2 and ESMFold can accurately predict the structures of *de novo* designed proteins from a single sequence, although only ESMFold was specifically trained for that purpose. Remarkably, the structures of complex multi-span membrane proteins with a water-accessible transmembrane channel could be predicted that way, although membrane proteins represent less than 2 % of the structures in the PDB^40^. Using designs generated on accurately assembled (“designable”) β-barrel backbones and designs incorporating non-natural backbone features (“un-designable”), we show that ESMFold could be used to rapidly and accurately assess backbone designability. It can predict the structure of a small protein in less than one minute, which makes it suitable for high-throughput analysis of large sets of sequences at the earliest stage of *de novo* design. However, once appropriate backbone constraints have been defined, only AlphaFold2 could capture the subtle differences resulting in proper folding or misfolding of the designed proteins. We show that it achieves better discriminative power than classic *ab initio* prediction for β-sheet proteins (which are notoriously difficult to design and fold *in silico*). Our results suggest that AlphaFold2 (but not ESMFold) can be used to prioritize designs for experimental validation. However, users should exercise caution since the prediction parameters might have to be adjusted to account for biophysical specificities of the designed fold, as demonstrated here with TMBs. By pre-filtering the TMB sequences based on their secondary structure and aggregation propensities and increasing the number of AlphaFold2 recycles, we designed a validation pipeline that could improve the success rate of *de novo* TMB designs 2.5 times. Moreover, we propose a new ESMFold prediction perturbation strategy that could be used to reveal differences in the robustness of intramolecular contacts in designed sequences and provide novel insight on the determinants of design success. We anticipate that DL-based structure prediction will open the door to the design of increasingly complex water-soluble and membrane proteins that were previously out of reach for *de novo* design.

Finally, this study suggests a potential risk in using ESMFold for protein design. In the examples presented here, even sparse contacts from LM (as much as 90 % of the sequence is masked) can be captured and amplified by the SM to return the correct structure. While desirable for structure prediction, such behavior might be problematic for protein design with an inverted ESMFold network. The iterative optimization of the protein sequence is guided by a loss function that defines a gradient towards a high-quality predicted structure. If a high-quality prediction is achieved faster than strong contacts are introduced, the designs will contain minimal information to trigger the prediction of the correct structure but will be unlikely to fold experimentally. Moving forward, a strategy that randomly masks input positions during design may help fixing this issue and designing robust proteins with good “melting temperature”.

## Material and methods

### AlphaFold modeling

AlphaFold’s Colabfold version (AlphaFold2.ipynb) was used through this study^44^. All predictions were made with the single_sequence MSA setting, no AMBER force-field relaxation, and no template. The number of recycles was set to 3 (for the designable/un-designable backbones test and for water-soluble β-barrels) or 48 (for transmembrane β-barrels). Five models were generated per input. The water-soluble β-barrels dataset was truncated to remove designs stabilized with disulfide bonds. Such designs usually differ only by a few mutations from a disulfide-free parent in the dataset but might have quite different experimental behavior. The transmembrane β-barrels dataset was truncated to exclude sequences with high β-sheet and/or aggregation propensities, as described in the main text.

### ESMFold modeling

ESMFold implemented in Colabfold (ESMFold_advanced.ipynb) was used through this study to generate structure predictions and visualize contacts from the language and the structure models. The number of recycles was set to 0.

### Evaluation metrics

The generated models were aligned to the coordinates of the original design model using TM-align^45^ to calculate the Root Mean Square Deviation (RMSD) and the TM-score. To analyze the ESMFold predictions, the pLDDT scores of the backbone heavy atoms (N, CA, O, C, CB) were averaged over the entire sequence. After confirming the skewness of the distributions with a normality test, the Mann Whitney U test was used to compare the distributions of confidence scores (pLDDT) and RMSD between populations of designs. The results were confirmed using a permutation test. When the data were represented in a box plot, the center line shows the median; the box limits show the first and third quartiles; the whiskers show the minimum and maximum of the distribution; the points show outliers.

### In silico melting with ESMFold

The analysis was done using a custom ESMfold colab notebook (https://colab.research.google.com/github/vorobieva/ColabFold/blob/main/beta/ESMFold_me

lting.ipynb). For each input sequence, perturbation was incrementally added to the prediction by applying a mask on randomly selected 30-95 % of residues in the sequence (5 % increment). For each perturbation step, sixty-four models were generated with different and randomly applied masking patterns of non-sequential positions in the sequence. The success rate at each step was defined as the fraction of models out of sixty-four with a predicted TM-score (pTM) higher than 0.75. To quantitatively compare the robustness of the predictions, the success rate as a function of the level of perturbation was fitted to a sigmoidal curve to calculate the inflection point of the transition.

## Acknowledgments

This work was supported by VIB core funding and by a FEBS excellence award to AAV. SO was supported by NIH DP5OD026389, NSF MCB2032259.

## Author contributions

AAV conceived the study with advice from SO. SO and AAV contributed code. AAV assembled input data. AM, CB and AAV performed the experiments and analyzed data. AAV wrote the manuscript with input from all authors.

## Data availability statement

All source data are available with this paper. These data, as well as the models generated for this study, are available on Zenodo: https://doi.org/10.5281/zenodo.8075911.

## Compute code statement

The code used to generate the predictions is available through the ColabFold repository: https://github.com/sokrypton/ColabFold and https://github.com/vorobieva/ColabFold//tree/main/beta/ESMFold_melting.ipynb to generate ESMFold melting predictions.

The code for processing, visualizing and analyzing results is available at: https://github.com/vorobieva/validate_de_novo_designed_proteins.

## References

1. Huang, P.-S., Boyken, S. E. & Baker, D. The coming of age of de novo protein design. Nature 537, 320–327 (2016).

2. Korendovych, I. V. & DeGrado, W. F. De novo protein design, a retrospective. Q. Rev. Biophys. 53, E3 (2020).

3. Kuhlman, B. et al. Design of a novel globular protein fold with atomic-level accuracy. Science 302, 1364–1368 (2003).

4. Brunette, T. et al. Exploring the repeat protein universe through computational protein design. Nature 528, 580–584 (2015).

5. Watson, J. L. et al. De novo design of protein structure and function with RFdiffusion. Nature (2023) doi:10.1038/s41586-023-06415-8.

6. Joh, N. H. et al. De novo design of a transmembrane Zn^2+^-transporting four-helix bundle. Science 346, 1520–1524 (2014).

7. Dou, J. et al. De novo design of a fluorescence-activating β-barrel. Nature 561, 485–491 (2018).

8. Kuhlman, B. & Bradley, P. Advances in protein structure prediction and design. Nat. Rev. Mol. Cell Biol. 20, 681–697 (2019).

9. Koga, N. et al. Role of backbone strain in de novo design of complex α/β protein structures. Nat. Commun. 12, 3921 (2021).

10. Rocklin, G. J. et al. Global analysis of protein folding using massively parallel design, synthesis, and testing. Science 357, 168–175 (2017).

11. Chidyausiku, T. M. et al. De novo design of immunoglobulin-like domains. Nat. Commun. 13, 5661 (2022).

12. Duran, A. M. & Meiler, J. Computational design of membrane proteins using RosettaMembrane. Protein Sci. 27, 341–355 (2018).

13. Vorobieva, A. A. Principles and methods in computational membrane protein design. J. Mol. Biol. 433, 167154 (2021).

14. Huang, P.-S. et al. High thermodynamic stability of parametrically designed helical bundles. Science 346, 481–485 (2014).

15. Bradley, P., Misura, K. M. S. & Baker, D. Toward high-resolution de novo structure prediction for small proteins. Science 309, 1868–1871 (2005).

16. Koga, N. et al. Principles for designing ideal protein structures. Nature 491, 222–227 (2012).

17. Leman, J. K. et al. Macromolecular modeling and design in Rosetta: recent methods and frameworks. Nat. Methods 17, 665–680 (2020).

18. Pan, X. & Kortemme, T. Recent advances in de novo protein design: Principles, methods, and applications. J. Biol. Chem. 296, 100558 (2021).

19. Barth, P., Schonbrun, J. & Baker, D. Toward high-resolution prediction and design of transmembrane helical protein structures. Proc. Natl. Acad. Sci. U. S. A. 104, 15682–15687 (2007).

20. Alford, R. F., Fleming, P. J., Fleming, K. G. & Gray, J. J. Protein structure prediction and design in a biologically realistic implicit membrane. Biophys. J. 120, 2042–2055 (2020).

21. Elazar, A. et al. De novo-designed transmembrane domains tune engineered receptor functions. Elife 11, p(2022).

22. Scott, A. J. et al. Constructing ion channels from water-soluble α-helical barrels. Nat. Chem. 13, 643–650 (2021).

23. Lu, P. et al. Accurate computational design of multipass transmembrane proteins. Science 359, 1042–1046 (2018).

24. Jumper, J. et al. Highly accurate protein structure prediction with AlphaFold. Nature 596, 583–589 (2021).

25. Baek, M. et al. Accurate prediction of protein structures and interactions using a three-track neural network. Science 373, 871–876 (2021).

26. Lin, Z. et al. Evolutionary-scale prediction of atomic-level protein structure with a language model. Science 379, 1123–1130 (2023).

27. Wu, R. et al. High-resolution de novo structure prediction from primary sequence. Preprint at https://www.biorxiv.org/content/10.1101/2022.07.21.500999v1 (2022).

28. Chowdhury, R. et al. Single-sequence protein structure prediction using a language model and deep learning. Nat. Biotechnol. 40, 1617–1623 (2022).

29. Anishchenko, I. et al. De novo protein design by deep network hallucination. Nature 600, 547–552 (2021).

30. Verkuil, R. et al. Language models generalize beyond natural proteins. Preprint at https://www.biorxiv.org/content/10.1101/2022.12.21.521521v1 (2022).

31. Goverde, C. A., Wolf, B., Khakzad, H., Rosset, S. & Correia, B. E. De novo protein design by inversion of the AlphaFold structure prediction network. Protein Sci. 32, e4653 (2023).

32. Novotný, J., Bruccoleri, R. E. & Newell. J. Twisted hyperboloid (strophoid) as a model of beta-barrels in proteins. J. Mol. Biol. 177, 567–573 (1984).

33. Murzin, A. G., Lesk, A. M. & Chothia, C. Principles determining the structure of β-sheet barrels in proteins I. A theoretical analysis. J. Mol. Biol. 236, 1369–1381 (1994).

34. Dhar, R., Feehan, R. & Slusky, J. S. G. Membrane barrels are taller, fatter, inside-out soluble barrels. J. Phys. Chem. B 125, 3622–3628 (2021).

35. Vorobieva, A. A. et al. De novo design of transmembrane β barrels. Science 371, p(2021).

36. Alford, R. F. et al. An Integrated Framework Advancing Membrane Protein Modeling and Design. PLoS Comput. Biol. 11, e1004398 (2015).

37. Dauparas, J. et al. Robust deep learning-based protein sequence design using ProteinMPNN. Science 378, 49–56 (2022).

38. Wang, J. et al. Scaffolding protein functional sites using deep learning. Science 377, 387–394 (2022).

39. Wicky, B. I. M. et al. Hallucinating symmetric protein assemblies. Science 378, 56–61 (2022).

40. White, S. H. Biophysical dissection of membrane proteins. Nature 459, 344–346 (2009).

41. Wang, S., Li, W., Liu, S. & Xu, J. RaptorX-Property: a web server for protein structure property prediction. Nucleic Acids Res. 44, W430–5 (2016).

42. Fernandez-Escamilla, A.-M., Rousseau, F., Schymkowitz, J. & Serrano, L. Prediction of sequence-dependent and mutational effects on the aggregation of peptides and proteins. Nat. Biotechnol. 22, 1302–1306 (2004).

43. Lin, Z. et al. Evolutionary-scale prediction of atomic-level protein structure with a language model. Science 379, 1123–1130 (2023).

44. Mirdita, M. et al. ColabFold: making protein folding accessible to all. Nat. Methods 19, 679–682 (2022).

45. Zhang, Y. & Skolnick, J. TM-align: a protein structure alignment algorithm based on the TM-score. Nucleic Acids Res. 33, 2302–2309 (2005).

