## Supplementary figures for "Validation of *de novo* designed water-soluble and transmembrane proteins by *in silico* folding and melting"

### Supplementary information

#### Supplementary figures

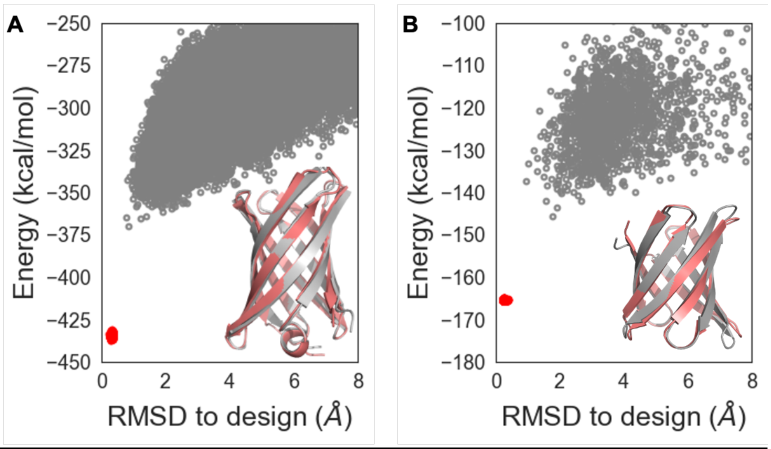


*Figure S1: Ab initio structure predictions of sequences generated based on designable (left) and undesignable (right) β-barrel backbones converge to low-energy structures similar to the design models.*

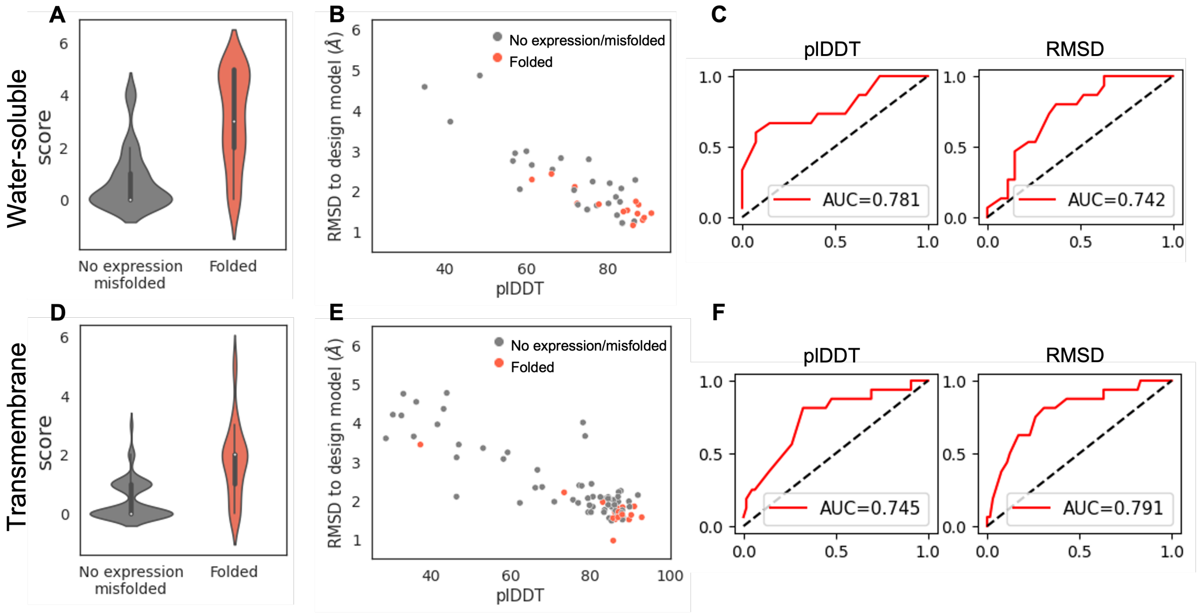


*Figure S2: (a, d) Out of five models, AlphaFold2 generates a higher number of high-quality models per input (pLDDT > 85.0 and RMSD < 1.75 Å) for folded than failed designs. (b, e) The lowest RMSD model can be used to select the most promizing designs. (c, f) Receiver-Operator Characteristic (ROC) curves quantifying the discriminatory power of AlphaFold2 plDDT and RMSD metrics to separate successful designs from unsuccessful designs. The True Positive Rate is plotted on the y-axis and the False Positive Rate on the x-axis. For each plot, the Area Under the Curve (AUC) is shown.*

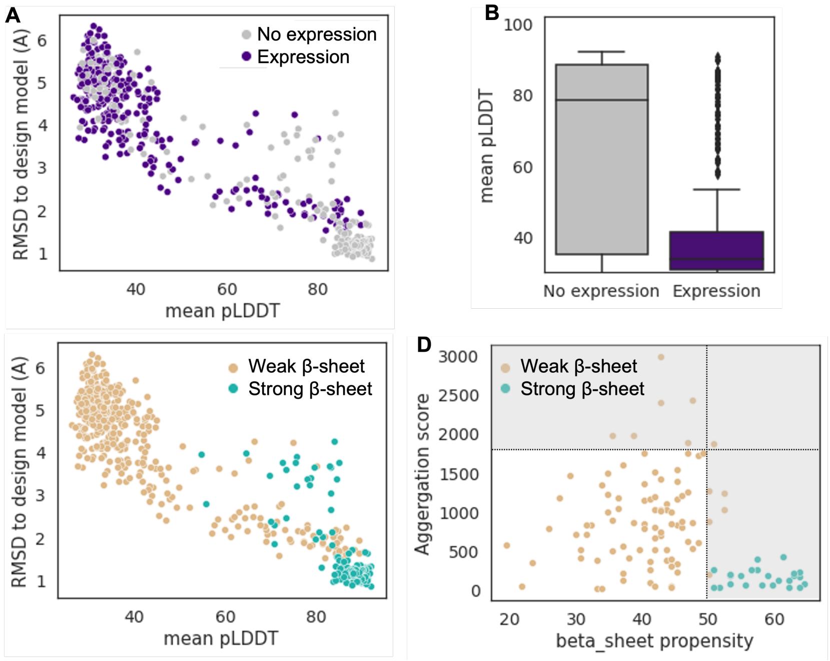


*Figure S3: (a-b) Standard AlphaFold2 pipeline (3 recycles) generates predicted structures attributed higher pLDDT, lower RMSD for TMB designs that failed to express experimentally. (c) AlphaFold2 predictions of TMBs designed with high propensity to form β-sheet (strong β-sheet) sample lower RMSD and higher pLDDT values than the sequences generated with negative designs (weak β-sheet). (d) TMB sequence are filtered to focus on the space accessible to experimental validation by applying empirical cutoffs on β-sheet and aggregation propensities.*

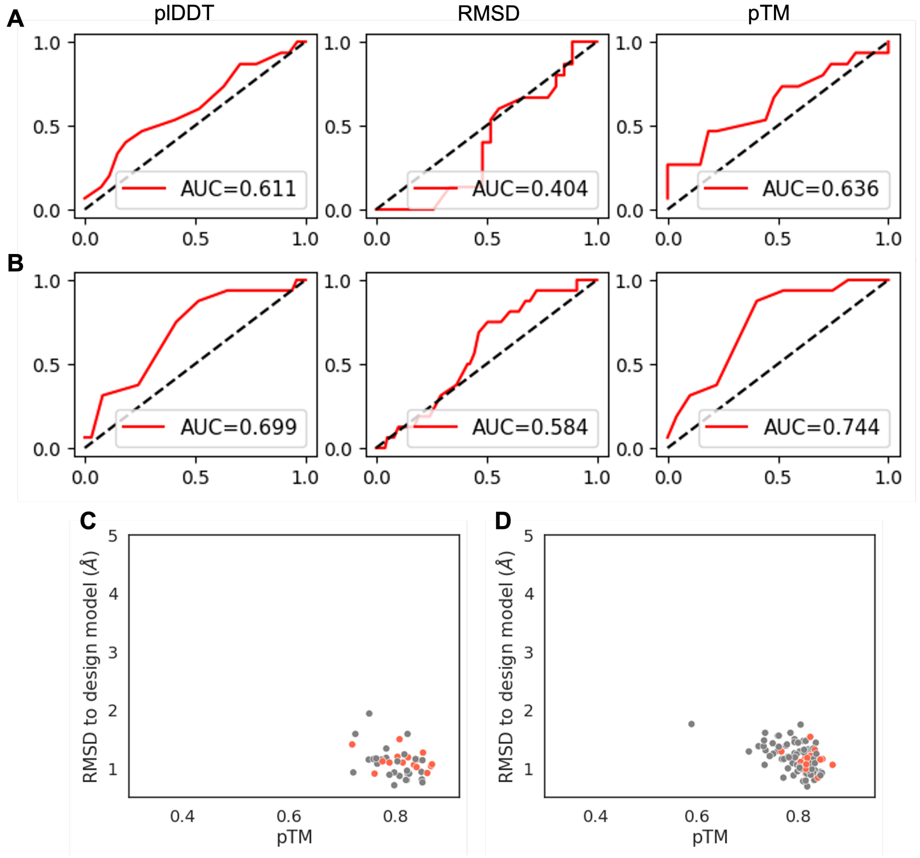


*Figure S4: (a, b) Receiver-Operator Characteristic (ROC) curves quantifying the discriminatory power of ESMFold plDDT, RMSD and predicted TMscore (pTM) metrics to separate successful from unsuccessful water-soluble (a) and transmembrane (b) β-barrel designs. The True Positive Rate is plotted on the y-axis and the False Positive Rate on the x-axis. For each plot, the Area Under the Curve (AUC) is shown. (c, d) pTM scores attributed to ESMFold predictions for de novo designed water-soluble (c) and transmembrane (d) β-barrel sequences.*
